# Long-term functional regeneration of radiation-damaged salivary glands through delivery of a neurogenic hydrogel

**DOI:** 10.1101/2022.05.19.491203

**Authors:** Jianlong Li, Sonia Sudiwala, Lionel Berthoin, Alison J. May, Seayar Mohabbat, Hanan Sinada, Eliza A. Gaylord, Noel Cruz Pacheco, Isabelle M.A. Lombaert, Oju Jeon, Eben Alsberg, Chelsea S. Bahney, Sarah M. Knox

## Abstract

Salivary gland acinar cells are severely depleted after radiotherapy for head and neck cancer, leading to loss of saliva and extensive oro-digestive complications. With no regenerative therapies available, organ dysfunction is irreversible. Here using the adult murine system, we demonstrate radiation-damaged salivary glands can be functionally regenerated via sustained delivery of the neurogenic muscarinic receptor agonist, cevimeline. We show that endogenous gland repair coincides with increased nerve activity and acinar cell division that is limited to the first week post-radiation, with extensive acinar cell degeneration, dysfunction and cholinergic denervation occurring thereafter. However, we discovered that mimicking cholinergic muscarinic input via sustained local delivery of a cevimeline-alginate hydrogel was sufficient to regenerate innervated acini and retain physiological saliva secretion at non-irradiated levels over the long-term (> 3 months). Thus, we reveal a novel regenerative approach for restoring epithelial organ structure and function that has significant implications for human patients.

**Teaser:** Novel application of an injectable neurogenic-based hydrogel for restoring the structure and function of radiation-damaged tissue.

## Introduction

Despite substantial improvements in the targeted delivery of ionizing radiation (IR) for the treatment of tumors, numerous neighboring organs, including the skin (*1*), heart (*2*) and salivary glands (*3*), are inadvertently damaged by off target effects. With IR being the primary treatment option for the elimination of head and neck cancers (67,000/yr in USA (*4*)), salivary glands represent one of the organ systems routinely injured, often resulting in permanent hyposalivation and related dry mouth condition xerostomia (*5*). As saliva plays a multitude of roles in maintaining oral and gastrointestinal health, pathological injury to these glands causes a diverse array of complications, including poor digestion, accelerated dental caries, periodontal disease, esophageal infections, and gastro-esophageal reflux disease (*5*). With no curative therapies available, these outcomes are irreversible.

Loss of saliva production after IR therapy has been primarily attributed to the destruction of the major secretory cell type, the acinar cell (*6, 7*). Both serous and mucous acini belonging to the 3 major salivary glands (i.e., parotid, submandibular and sublingual) secrete a fluid containing water, electrolytes, mucus and proteins e.g. enzymes and immunoglobulins, into the ductal system for deposition within the oral cavity. Numerous studies have demonstrated these cells to be gradually destroyed after IR exposure, with rodents and humans exhibiting extensive loss of serous acinar cells within 2-3 months (*7, 8*). Unfortunately, no regenerative treatments are currently available to promote the repopulation of acinar cells and repair organ function post-IR. Furthermore, current palliative treatment options, such as salivary stimulants, topical agents, saliva substitutes, systemic sialagogues and mouth rinses, do not recapitulate the multi-functional roles of saliva, thus resulting in pronounced patient morbidity over their lifetime (*9*).

In addition to destroying salivary gland acinar cells, IR therapy has also been shown to deplete the parasympathetic nerve supply (*10*). Parasympathetic nerves are an essential driver of acinar cell function, maintenance and repair, with studies over the last 60+ years showing a clear requirement for these nerves in glandular function and homeostasis (*10*–*12*), with nerve damage resulting in glandular atrophy (*11, 12*) and re-innervation promoting tissue regeneration (*10, 13, 14*). Multiple studies have further revealed that the positive feedback loop between parasympathetic nerves and acinar cells is crucial to homeostasis, and that loss in either signaling loop e.g., nerve acetylcholine-muscarinic signaling to acini or acinar neurotrophic signaling to nerves, results in tissue degeneration (*15*). More specifically, in vivo studies indicate that stimulation of muscarinic receptors is necessary to promote the division of acinar (SOX2^+^) progenitors and their (SOX2^-^) transit-amplifying progeny (*10*), supporting a role for this pathway in acinar cell replenishment. Furthermore, application of a synthetic muscarinic agonist (e.g., carbachol) to ex vivo cultures of irradiated adult tissue is sufficient to repopulate acinar cells at least during the first 48 hours (h) following IR treatment (*10*). However, whether acinar cells retain their ability to undergo cell division in response to muscarinic activation (i.e., beyond 48 h post-IR), and if re-establishing the muscarinic pathway post-IR can regenerate functional acini, remains unknown. Moreover, whether parasympathetic nerve-acinar cell communication can be restored in the irradiated tissue to mediate long-term organ function and homeostasis has not been determined.

*Through a series of in vivo studies*, we show that local delivery of the muscarinic receptor (CHRM1 and 3) agonist, cevimeline, encapsulated in an alginate hydrogel (ALG) to irradiated salivary glands is highly efficacious in maintaining saliva secretion and glandular function over an extensive time period that is typically associated with glandular degeneration. We demonstrate that radiation-induced loss of acinar cell proliferation and parasympathetic innervation is rescued by muscarinic receptor activation, thereby ensuring the preservation of the acinar cell pool and a functional nerve supply required for homeostatic and secretory processes. Thus, we provide a novel therapeutic approach for the long-term restoration of salivary gland function after radiation therapy.

## Results

### Acinar cells undergo a time limited regenerative response 1 week following radiation exposure that correlates with an increase in neuronal-related genes

We previously demonstrated that mucous acinar cells in the murine salivary (sublingual) gland are actively replaced by SOX2^+^ progenitor cells during the first 2 weeks following IR treatment (*10*). However, whether acinar cells continuously repopulate the organ during this time frame, and if changes in nerve-derived muscarinic input are involved, remains to be determined. Thus, we first questioned the timing of endogenous regenerative events in response to a single 10 Gy dose of IR by sequencing the transcriptomes of glands from adult (6-7 wks) mice at 7- and 14-days post-IR and comparing to non-IR controls (**Fig.1A, Fig.S1A**). Differential gene expression analysis (DESeq2 (*16*); *P*adj<0.1; 1.5-fold change cutoff) revealed extensive alterations in gene expression at the 7-days post-IR time point compared to non-IR controls, with 1354 genes being upregulated and 1122 downregulated (**Fig.1B, Fig.S1B, Data S1**). Gene ontology (GO) enrichment analysis of the significantly changed genes using g:Profiler (*17*) identified a striking increase in genes associated with cell division (**Fig.1A, B**). These included positive regulators of cell cycle progression (e.g., *Cdk1, Bub1, Ccnd1, Plk1;p<*0.01), with particular enrichment for genes associated with mitotic events, such as spindle assembly, chromatid segregation (e.g., *Tpx2, Nek2, Kif23, Birc5;p<*0.01) and cytokinesis (e.g., *Cep55*; *Ndc80;p<*0.001) (**Fig.1A**). The downregulated genes were primarily involved in protein synthesis and mitochondrial oxidative phosphorylation (**Fig.S1B**), suggesting a reduction in secretory processes occurs simultaneously with an increase in proliferation over the first 7 days.

To determine changes in gene expression across time, we next measured alterations in gene transcript levels at 14-days post-IR compared to the 7-day time point and found 323 genes to be upregulated and 324 downregulated (**Data S2)**. As shown in **Figure 1**, the most significant increase was associated with the immune system (128 genes), with extensive upregulation of genes/pathways involved in the innate immune response that is indicative of tissue injury, including the master transcription factors *Irf5* (*p*<0.05) (*18*) and *Irf7* (*p*<0.01) (*19*), and markers of systemically recruited monocytes (e.g., *Ccr2 and Lyz2; p<*0.01) (*20*) (**Fig.1A**). Together with previous findings in irradiated submandibular salivary glands showing that edema or immune cell infiltrate does not occur during the first 10 days following IR (*21*), this outcome suggests that immune cell recruitment occurs between 10-14 days post-IR. The largest cohort of downregulated genes at 14-days (101 genes) was associated with the cell cycle (**Fig.1A, B**), with many of the same genes that had increased at 7-days post-IR, including *Bub1, Ccnd1, Plk1, Tpx2, Kif23, Birc5 and Ndc80* (*p*<0.05), returning to non-IR levels. In addition to the decrease in genes involved in mitosis, those involved in cell cycle progression were also downregulated (e.g., *Rb1, Usp2, Cdk1, Ccne1, Ccnb1; p*<0.05), an outcome suggestive of cell cycle exit by day 14 post-IR (**Fig.S1B)**. Finally, we determined whether the 14-days post-IR tissue was returning to the non-IR state by comparing it to non-IR controls. In contrast to 7-day post-IR glands, the number of genes differentially expressed at 14 days was low, with only 308 genes being significantly upregulated and 121 downregulated (**Fig.S1C and Data S3**). Analysis of GO terms revealed an upregulation of genes involved in immune response, while only a few genes associated with responses to ER stress and negative regulation of cellular metabolisms were downregulated. Together, these results indicate that salivary glands at 7-days post-IR are in a phase of increased cell turnover, but that by 14-days post-IR cell proliferation is reduced to basal levels while there is an increasing tissue immune response.

**Fig. 1.**
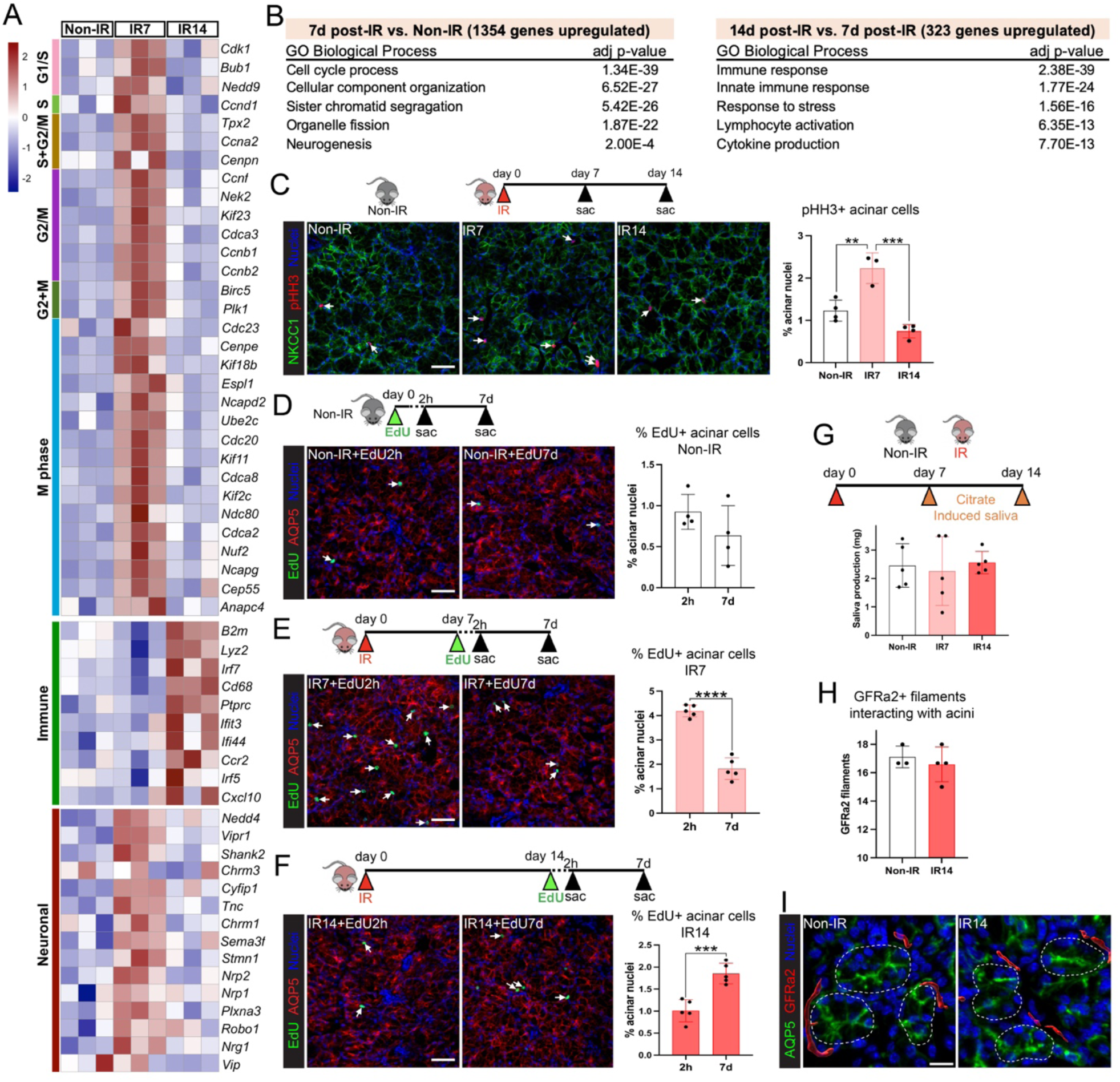
Acinar cells undergo a time limited regenerative response 1 week following radiation exposure that correlates with increased neuronal activity. **A**, Heatmap showing mitotic, immune and neuronal gene expression in sublingual salivary glands at 7- or 14-days post-IR (single 10 Gy dose), compared to non-IR controls, n=3 mice per group. **B**, Gene Ontology (GO) analysis of significantly upregulated genes (FC>1.5, p<0.05). **C**, Sublingual glands from non-IR (control), 7- and 14-days post-IR mice were immunostained for the mitotic marker phospho-histone H3 (pHH3, red) and acinar marker, Na-K-Cl cotransporter (NKCC1, green), nuclei in blue (Hoechst 33342). The number of pHH3^+^ acinar cells was quantified and expressed as a percentage of the total number of acinar nuclei (graph). White arrows point to pHH3^+^ NKCC1^+^ proliferating acinar cells. **D-F**, Immunofluorescent detection of EdU^+^ cells (green) in sublingual glands at 2 h and 7 days following a single injection of EdU into non-IR mice (**D**), and mice at 7-(**E**) and 14-days post-IR (**F**), with white arrows pointing to EdU^+^ AQP5^+^ acinar cells. The number of EdU^+^ acinar cells was quantified for each treatment and is expressed as a percentage of total acinar cell number (graphs). Acinar cells were determined by immunostaining of aquaporin 5 (AQP5, red), nuclei in blue (Hoechst 33342). **G**, Citrate-induced saliva collection at 7- and 14-days post-IR, non-IR as control. **H-I**, Quantification (**H**) of parasympathetic nerves (GFRa2, red) innervating acini (AQP5, green) in sublingual glands at 14 days post-IR vs. non-IR control. Nuclei are labeled using Hoechst 33342 (blue). Scale bars = 40 μm in **C-F** and 15 μm in **I**. Each dot in the bar graph represents a biological replicate. Data are expressed as mean ± s.d. \**P*<0.05; \*\**P*<0.01; \*\*\**P*<0.001; \*\*\*\**P*<0.0001; an unpaired two tailed t-test for two-group comparisons was applied to graphs D-F, H; a one-way analysis of variance followed by a Dunnett’s test for multiple comparisons was applied to graphs C and G.

Next, we performed immunofluorescent analysis to confirm changes in cell turnover and identify the proliferating cell type(s) during the 2-week time period post-IR. First, epithelial cell proliferation was quantified by immunostaining tissue for phospho-histone H3 (pHH3), a marker of the M phase of the cell cycle (*22*), and the acinar cell marker Na-K-Cl cotransporter (NKCC1). In line with the abundance of cell cycle-related genes, glands at 7-days post-IR exhibited an 81 % increase in pHH3^+^ mitotic cells, restricted to the acinar cell population, compared to the non-IR controls (*P*<0.01, **Fig.1C and Fig.S1D**). Consistent with the transcriptomic analysis, acinar cell proliferation at 14-days post-IR was significantly reduced compared to 7-days (a 67% reduction; p<0.001, **Fig.1C and Fig.S1D**) but was not significantly different from non-IR controls (**Fig.1C and Fig.S1D**).

To verify that acinar cells were undergoing cell turnover, we performed a pulse-chase assay in which EdU (5-ethynyl-2′-deoxyuridine) was systemically injected into non-IR, or irradiated mice at 7- or 14-days post-IR. The number of EdU^+^ acinar cells (as a percentage of total acinar cells) was then quantified after a 2 h or 7-day chase at each of these time points e.g., day 7 +2 h or day 7 +7 days. Consistent with previous studies (*23*), acinar cell turnover under non-IR (homeostatic) conditions was low, as shown by the number of EdU^+^ acinar cells after the 2 h chase being similar to that after 7 days (**Fig.1D and Fig.S1E**). In contrast, we found extensive acinar cell proliferation at 7 days post-IR (day 7 +2 h), with a 4-fold increase (p<0.001) in the number of EdU+ cells at this time point compared to non-IR controls (**Fig.1E and Fig.S1F**). Moreover, these cells underwent substantial turnover, as demonstrated by the significant reduction in EdU+ acinar cells after the 7-day chase (>2-fold decrease; *p*<0.0001) (**Fig.1E and Fig.S1F**). In comparison, active acinar cell turnover at 14-days post-IR was greatly diminished. Although the 2 h chase suggested 14-days post-IR glands had similar levels of cell division to that of homeostatic non-IR glands (**Fig.1F and Fig.S1G**), we found an 84% increase (*p*<0.001) in EdU^+^ acinar cells after the 7-day chase, suggesting originally EdU-incorporated acinar cells had undergone a round of cell replication but did not undergo further cell division (**Fig.1F and Fig.S1G**). Thus, these results are consistent with both our pHH3^+^ quantification and transcriptomic analysis and confirm that salivary glands undergo an early phase of active acinar cell proliferation and replacement in response to IR that is lost by 14 days.

Given the known role of parasympathetic nerves in salivary gland secretory function and acinar cell replacement (*10, 24*), we next asked whether the functional nerve supply to acini was altered at 7- or 14-days post-IR compared to non-IR controls. Bulk RNA-seq analysis identified a significant upregulation of genes involved in neuronal patterning, nerve migration and neurogenesis (*Nrg1, Nedd4, Nrp1, Nrp2, Robo1, Cyfip1*) (*25, 26*), synapse assembly/activity (*Stmn1*) (*27*) and nerve function (*Vip* (*10*)) at 7-days post-IR, suggesting radiation exposure activates neuronal programs at the same time as acinar cell replacement (**Fig.1A, B)**. Importantly, we also found the muscarinic receptor 1 (*Chrm1*), which upon activation promotes proliferative signaling in acinar progenitors and a subset of transit-amplifying cells (*28*), to be upregulated (**Fig.1A)**, an outcome suggestive of an increase in nerve-acinar cell communication at 7-days post-IR. However, by 14-days post-IR, neuronal genes were reduced, with levels returning to those of non-IR controls, indicating a return to homeostatic innervation.

Next, we analyzed changes in nerve-acinar cell function by measuring physiological saliva output at 7-and 14-days post-IR through induction of the gustatory-salivatory reflex (*29*). The gustatory-salivatory reflex involves afferent input from the tongue to the dorsal medulla which synapse with parasympathetic neurons innervating the salivary gland, with stimulation of these nerves driving saliva secretion (*30, 31*). Gustatory nerves were stimulated through the local application of citric acid to the tongue, and saliva collected over 20 minutes, as performed previously (*31*). Saliva output from mice at both 7- and 14-days post-IR was similar to non-IR controls (**Fig.1G**), suggesting radiation does not damage parasympathetic nerve-acinar cell interactions or nerve function at these time points. This was further confirmed by immunofluorescent analysis, followed by surface rendering using Imaris image analysis software, revealing similar levels of GFRa2^+^ parasympathetic nerves in close proximity to acini at 14-days post-IR and in non-IR controls (**Fig.1H, I**). However, similar to previous findings (*32*), direct stimulation of saliva secretion from acinar cells through systemic injection of the synthetic global muscarinic agonist (can activate *Chrm1-5*), pilocarpine, revealed a 20% reduction in maximal saliva output (totaling all 3 major salivary glands) at 14-days, but not 7-days, post-IR (*p*<0.01, **Fig.S1H**), confirming previously described, specific pilocarpine-stimulated secretory dysfunction by 2-weeks post-IR as a result from damage to muscarinic receptor-coupled signal transduction in parotid acinar cells (*33*).

### Acinar cells remain receptive to muscarinic-induced proliferation 14-days post-IR, with local delivery of a cevimeline alginate hydrogel extending the time period of cell division

Using an ex vivo salivary gland (sublingual) organoid model, we previously showed that muscarinic receptor activation within the first 48 h following IR exposure results in the repopulation of acini by SOX2^+^ progenitor cells (*10*) (**Fig.2A**). However, whether irradiated acinar cells remain receptive to muscarinic receptor-induced proliferation weeks after IR exposure is unknown. To test the regenerative potential of acinar cells, we subcutaneously administered the muscarinic agonist cevimeline (CV) locally to the neck region, overlying both the submandibular and sublingual glands (**Fig.2B-C**), at 14-days post-IR. The 14-days post-IR timepoint was chosen based on the reductions seen in acinar cell turnover and pilocarpine-induced maximal saliva output, as compared to 7-days post-IR (**Fig.1F, G** and **Fig. S1E**). Given the identity of acinar progenitors is known in the sublingual, but not the submandibular gland, we first quantified the number in pHH3^+^SOX2^+^ acinar progenitor cells and their progeny, the pHH3^+^SOX2^-^ transit amplifying cells, in response to CV at 18 h, 3- and 7-days after administration (**Fig.2D**). As shown in **Figure 2E**, local delivery of free CV significantly induced acinar cell proliferation 18 h after administration, with a 1.8-fold increase in pHH3^+^SOX2^+^ progenitors (white arrows, cell number shown as percentage of total number of acinar nuclei), and a 4.5-fold increase in pHH3^+^SOX2^-^ transit amplifying acinar cells (white arrowheads) compared to saline-treated controls (**Fig.2E, Fig.S2A**). However, cell proliferation returned to baseline (saline) levels by 3-days post-CV administration (**Fig.2E**, CV), thereby demonstrating that acinar cells are still receptive to muscarinic-driven proliferation but that a single injection of free CV-mediated cell cycle entry is highly transitory.

**Fig. 2.**
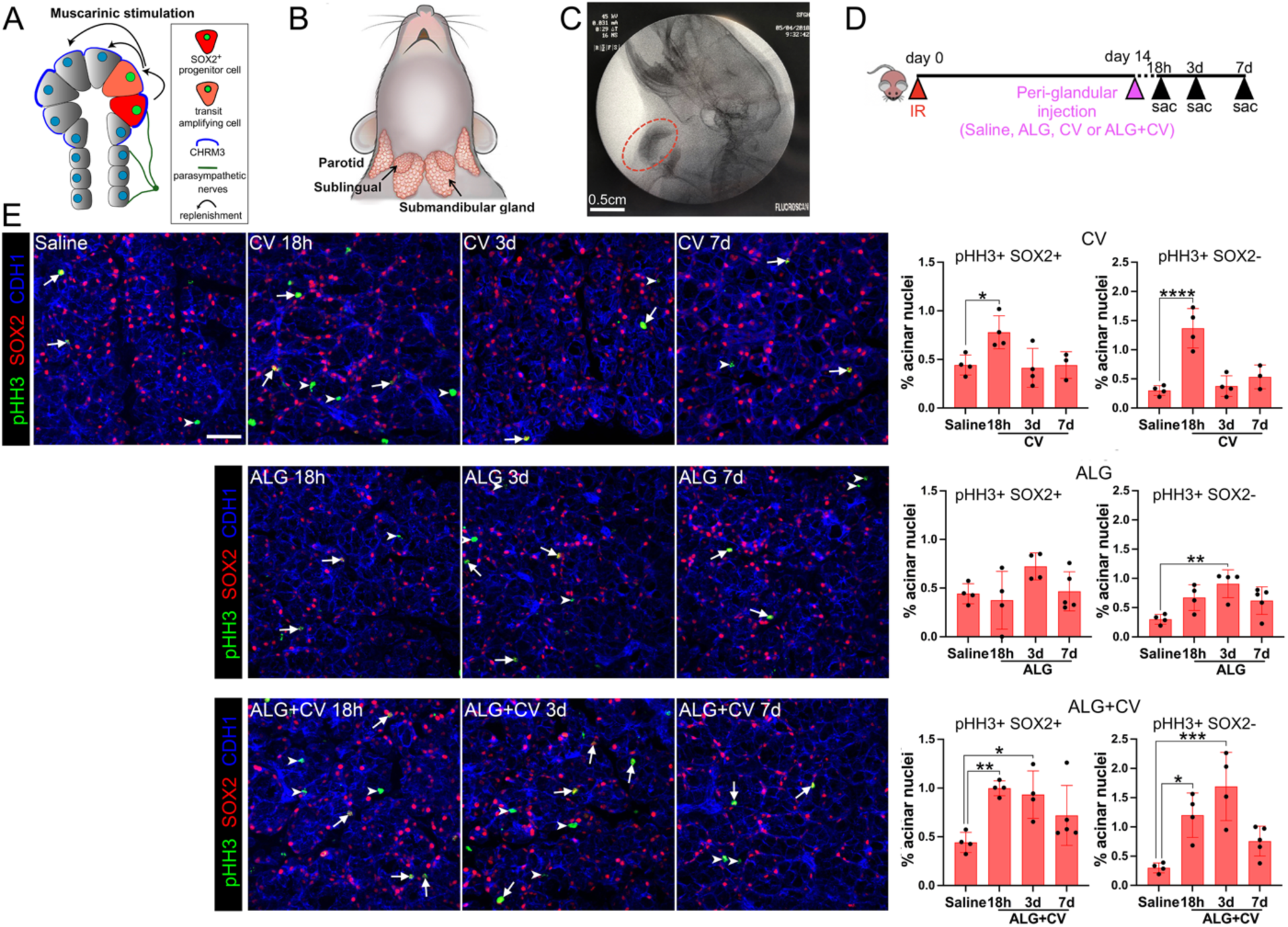
Cevimeline, free or incorporated into an alginate hydrogel (ALG), stimulates acinar cell replenishment 2 weeks post-radiation. **A**, Schematic showing salivary acinar replacement in healthy tissue is regulated by parasympathetic innervation. Upon muscarinic stimulation, SOX2+ progenitor cells divide and produce daughter acinar cells. **B**, Location of the major murine salivary glands - parotid, submandibular and sublingual. **C**, ALG was delivered via injection into the area immediately adjacent to the salivary glands and imaged via X-ray (Fluroscan). ALG is marked by a red circle. **D**, Radiation and treatment regimen. Salivary glands were collected and analyzed 18 hours (h), 3 days (3d) and 7 days (7d) after treatments with saline, cevimeline (CV) or CV+ALG. **E**, Immunofluorescent analysis of SOX2^+^ acinar progenitors and SOX2^-^ transit amplifying acinar cells from different treatment conditions (shown in D) using antibodies to the mitotic marker, phospho-histone H3 (pHH3, green), SOX2 (red) and the epithelial marker, E-cadherin (CDH1, blue). Arrows point to pHH3^+^ SOX2^+^ cells, arrowheads point to pHH3^+^ SOX2^-^ cells. Scale bar = 40 μm. Dots in the bar graphs represent biological replicates. Data are expressed as mean ± s.d. \**P*<0.05; \*\**P*<0.01; \*\*\**P*<0.001; \*\*\*\**P*<0.0001; a one-way analysis of variance with Dunnett’s multiple comparison test was applied.

Given this result, we next asked whether slow release of CV via incorporation into an alginate hydrogel (ALG) could promote a longer period of mitotic activity compared to free CV. Alginate is a biocompatible, naturally occurring polymer which is considered safe for drug delivery (*34*), and has been used in various drug-release settings (*35*–*37*). Similar to pilocarpine (*37*), cevimeline forms non-covalent interactions with oxidized alginate, which allows its diffusion from the hydrogel once placed. Hereto, ALG or ALG+CV were injected subcutaneously and peri-glandular (sublingual and submandibular) at 14-days post-IR (**Fig.2C**), and cell proliferation was measured 18 h, 3- and 7-days later. As shown in **Figure 2E** (lower panels), a single dose of ALG+CV significantly increased both SOX2^+^ progenitor and SOX2^-^ transit amplifying cell proliferation at both 18 h (2-fold and 4-fold, respectively) and 3 days (∼2-fold and ∼5-fold, respectively) compared to saline-treated controls, with increased cell proliferation remaining at 7 days, albeit not significantly (**Fig. 2E**, ALG+CV). Notably, ALG by itself was also able to promote cell proliferation (**Fig.2E**, ALG), although this was only significantly increased in SOX2^-^ transit amplifying cells at 3 days post-injection (3-fold; p<0.01; **Fig.2E**, ALG graph). To ensure muscarinic agonist-induced acinar cell proliferation was not restricted to the sublingual gland, we also quantified pHH3^+^MUC10^+^ serous acinar cells in the submandibular gland (**Fig. S2B-D**) and found a significant increase in cell division at 18 h and 3-days post-injection of ALG+CV (2.4-fold (p<0.01) and 2.3-fold (p<0.05) increase, respectively), with the values returning to baseline by 7- days (**Fig. S2D**). Together, these data demonstrate that sublingual and submandibular acinar cells can still be induced to divide in vivo at 2 weeks post-IR through stimulation of muscarinic receptors, and that the effect of this mitotic agent can be extended up to at least 3-7 subsequent days through the incorporation of CV into an alginate hydrogel.

### A single-dose of ALG+CV transiently rescues saliva secretion and reduces glandular degeneration after radiation

Given the proliferative response of irradiated acinar cells to CV, we next asked whether a single dose of CV or ALG+CV at 14 days post-IR could maintain saliva secretion and salivary gland architecture over a 10-week time period. This 10-week time frame was chosen based on rodent models showing extensive salivary gland degeneration 8-12 weeks after radiation exposure (*7*). Mice were irradiated at day 0 and treated with ALG+CV, free CV, ALG only or saline (control) 14 days later, and physiological levels of citrate-driven (nerve-mediated) saliva output being measured each week through gustatory stimulation (**Fig.3A**). A maximal (pilocarpine-driven) saliva secretion assay was also performed at 68 days, and tissue was then extracted for immunofluorescent analysis at 70 days (10 weeks, **Fig.3A**).

As shown in **Figure 3B**, saline-treated IR (control) mice maintained physiological saliva secretion at non-IR levels until 41-days post-IR (6 weeks), when a significant decrease was measured (30%,p<0.05), an outcome sustained over the following weeks. In contrast, IR mice treated with free CV or ALG+CV were able to maintain secretory output at similar levels to the non-IR controls beyond 41 days post-IR (**Fig. 3B**). At 49-days post-IR, both CV and ALG+CV treated mice showed significantly increased saliva secretion compared to saline treated IR mice (87% and 78% increase, respectively, *p<*0.05). However, only ALG+CV treatment was able to significantly maintain saliva output at non-IR levels by day 56 (68% increase compared to saline IR controls, *p<*0.05) (**Fig.3B**), suggesting that slow release of CV promotes extended functional recovery. By day 63, however, saliva secretion from ALG+CV treated mice was similar to saline IR mice, indicating that physiological function was no longer preserved (**Fig.3B**). Similarly, maximal saliva secretion from all irradiated mice (saline control or treated) was reduced below non-IR controls at day 68 (**Fig.3C**). Notably, maximal saliva levels in irradiated mice at day 68 were not significantly different from those at 14 days post-IR, suggesting that the level of tissue damage at this later time point did not further reduce muscarinic receptor-mediated saliva output. Thus, while a single dose of ALG+CV and CV treatment maintains physiological saliva output up to 8 weeks post-IR, it is not capable of preserving saliva secretion long-term.

**Fig. 3.**
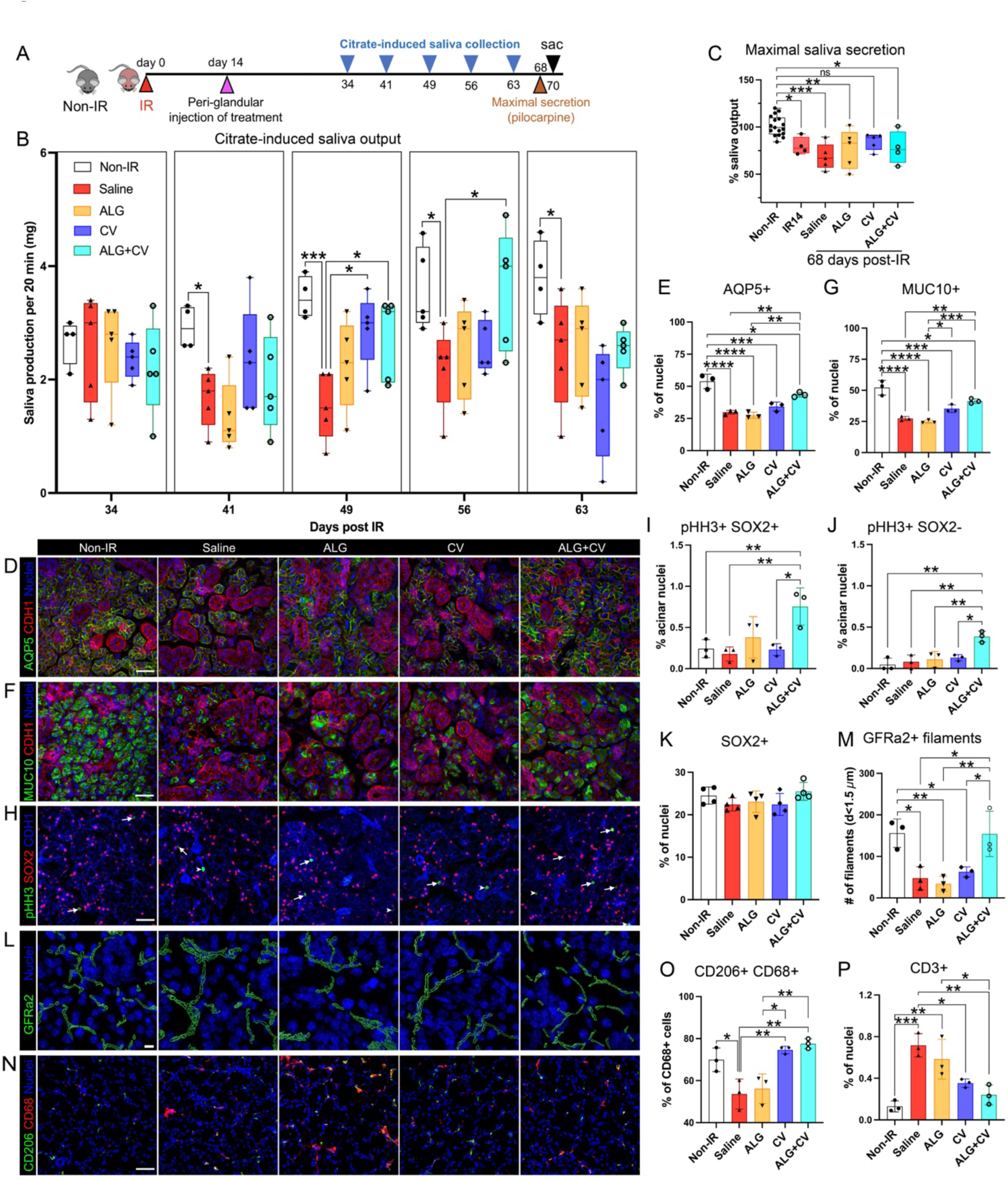
A single-dose of ALG+CV transiently rescues saliva secretion and reduces glandular degeneration after radiation. **A**, Regimen for radiation, treatment and saliva collections. **B**, Physiological saliva output levels were measured weekly from 34 to 63 days post-IR using gustatory stimulation (citric acid applied to tongue). **C**, Maximal saliva secretion in response to systemic pilocarpine administration at 68 days. Data was normalized to non-IR controls. **D-G**, Representative images and quantification of acinar cells expressing AQP5 (**D** and **E**) or the secreted protein MUC10 (**F** and **G**). E-cadherin (CDH1, red) marks all epithelial cells, Nuclei are stained with Hoechst 33342 (blue**). H-K**, Representative images (**H**) and quantification of mitotic (pHH3^+^, green) acinar progenitor cells (**I**, SOX2^+^, red) and transit amplifying cells (**J**, SOX2^-^) as well as total number of SOX2^+^ cells (**K**). **L-M**, Representative images (**L**) and quantification (**M**) of GFRa2^+^ parasympathetic nerve filaments (green). GFRa2^+^ filaments with diameters of less than 1.5 µm (innervate the acini) were quantified. **N-P**, Representative images of macrophages (**M**) and quantification of M2-polarized CD68^+^CD206^+^ macrophages (**O**) and CD3^+^ T cells (**P**). Scale bars in C, F, H and N = 40 µm, in L = 10 µm. Dots in the graphs represent biological replicates. Data are expressed as mean ± s.d. \**P*<0.05; \*\**P*<0.01, \*\*\**P*<0.001, \*\*\*\**P*<0.0001; a one-way analysis of variance with Dunnett’s multiple comparison test was applied for all graphs except B; a two-way analysis of variance with Dunnett’s multiple comparison test was applied to B.

We next proceeded to analyze salivary glands for alterations in tissue structure and function by measuring changes in secretory cell number and secreted protein synthesis. Given the submandibular gland is responsible for at least 60% of saliva secretion in rodents (*38*) and its serous acini are severely damaged by IR (*7*), we first focused on acinar cell architecture and protein biosynthesis within this gland type by immunostaining for the water channel aquaporin 5 (AQP5) and the secreted protein mucin 10 (MUC10). Consistent with the known degeneration of irradiated submandibular glands 8-12 weeks after IR exposure (*7*), saline-and ALG-treated tissue showed extensive destruction of functional acini at 70 days (10 weeks) post-IR (**Fig. S3A**), with a 50% decrease in both the number of AQP5^+^ (*P*<0.0001; **Fig.3D, E**) and MUC10^+^ acinar cells (*P*<0.0001; **Fig.3F, G**). In comparison to non-IR glands, CV-treated glands showed only a 37% reduction in AQP5^+^ acinar cells (*P*<0.001; **Fig.3D, E)** and a 33% decrease in MUC10^+^ acinar cells (*P*<0.001; **Fig.3F, G**), suggesting that the short (18 h) induction of cell proliferation with CV alone at 14 days post-IR (**Fig. S2D**) already positively impacted acinar cell number. However, preservation of secretory acini was even more pronounced in ALG+CV-treated glands, with only a 20% reduction (*P*<0.01) in the number of AQP5^+^ and MUC10^+^ acinar cells compared to non-IR glands (**Fig.3D-G**), suggesting that the longer-term induction of cell proliferation after delivery of ALG+CV at 14 days post-IR (**Fig. S2D**) significantly maintains acinar cells beyond the IR-induced degeneration phase (>8 weeks).

Given these outcomes, we next determined whether acinar progenitors remained in a regenerative state by probing for mitotic SOX2^+^ acinar progenitors and SOX2^-^ transit amplifying cells in the sublingual gland at 70 days post-IR. In addition to utilizing the known progenitor identities in the sublingual gland, mucous acini are less susceptible to IR-induced destruction compared to serous cells (*39, 40*), thus allowing for a critical analysis of total cell division without extensive cell loss. Strikingly, IR glands treated with ALG+CV, but not ALG or CV, showed a significant increase in dividing acinar cells, with a 5-fold increase in pHH3^+^SOX2^+^ progenitors (*P*<0.01) and a 10-fold increase in pHH3^+^SOX2^-^ transit amplifying (*P*<0.01) cells compared to saline treated IR controls (**Fig.3H-J**). However, despite ALG+CV promoting progenitor cell proliferation, the overall number of SOX2^+^ progenitors was unchanged compared to non-IR controls (**Fig.3K**), suggesting SOX2^+^ cells are likely activated to undergo asymmetric division. Thus, these data indicate that in spite of the reduction in salivary gland secretory function by 10 weeks, acinar cells continue to proliferate in irradiated glands in response to ALG+CV, resulting in improved acinar cell number and glandular integrity.

Given that parasympathetic nerves regulate physiological saliva secretion as well as acinar cell replacement (*10, 41*), we next determined whether CV-treatment preserved parasympathetic innervation of the acini. In the salivary glands, thick nerve bundles travel alongside ducts, and axons begin to separate (i.e., defasciculate) as they innervate individual acini. However, during injury, axons degenerate or retract (*42*), resulting in the appearance of thicker nerve bundles and reduced innervation of secretory cells. Therefore, to detect changes in neuronal presence, we quantified the number of thin (<1.5µm in diameter) GFRa2^+^ nerve filaments within the acinar compartment using Imaris surface rendering and filament tracing tools (**Fig.S3B**). Consistent with tissue injury, we measured a significant reduction in the number of thin GFRa2^+^ filaments innervating the acini of saline-(70% decrease; p<0.05) and ALG-treated mice (78% decrease; p<0.01) compared to non-IR controls (**Fig.3L**,**M and Fig.S3B**), indicating IR results in acinar cell denervation by this time point. Innervation of acini was also substantially reduced in CV-treated mice (*P*<0.05), with a 60% decrease in GFRa2^+^ axon filaments compared to non-IR tissue (**Fig.3L, M and Fig.S3B**). Strikingly, however, innervation of ALG+CV acini was similar to that of the non-IR controls (**Fig.3L,M and Fig.S3B**), suggesting that, despite the loss in physiological saliva secretion, nerve-acinar interactions remained intact. To ensure acinar cells within ALG+CV treated glands remained capable of receiving signals through the muscarinic pathway, we immunostained tissue at 70 days post-IR for the muscarinic receptor CHRM3 (*43*), the key driver of acinar cell secretion. We found CHRM3 levels in the acinar cells of ALG+CV-treated glands to be similar to non-IR cells (**Fig.S3C**), indicating that muscarinic receptor expression was maintained but that acinar cells had reduced secretory function in response to cholinergic signals.

Based on the fact that parasympathetic nerves can promote an anti-inflammatory environment (*44*), we next questioned whether cevimeline also impacted immune cell infiltration at 70 days. We found a drastic change in inflammatory cells in irradiated tissue treated with saline or ALG compared to non-IR controls (**Fig.3N-P and Fig.S3D-F**). With respect to the innate immune response, all irradiated tissues, regardless of treatment, demonstrated a significant increase in the total number of CD68^+^ macrophages (*P*<0.01, pan-macrophage marker) (**Fig.S3E-F**). However, the ratio of pro-reparatory M2 (CD68^+^ CD206^+^) to pro-inflammatory M1 (CD68^+^ CD206^-^) macrophages (*45*) was significantly elevated in ALG+CV and CV-treated tissues compared to saline-treated IR salivary glands (**Fig.3O**). As expected from previous studies (*46*), we found a 5-fold increase in the number of CD3^+^ T cells in saline and ALG treated mice, suggestive of activation of degenerative processes; p<0.001 for saline and p<0.01 for ALG; **Fig.3P and Fig.S3C**). However, mice treated with either ALG+CV or CV showed significantly fewer CD3^+^ T cells compared to saline IR controls (3-fold decrease; p<0.05 and 2-fold decrease; p<0.01, respectively) and were not significantly different from non-IR glands (**Fig.3P**). Together, these outcomes suggest that cevimeline promotes an anti-inflammatory/pro-reparatory environment.

In summary, despite saliva secretion not being fully restored, these data indicate that slow-release of ALG+CV preserves the acinar compartment through promoting proliferation and maintaining the surrounding nerve supply while activating pro-repair immune responses.

### Multiple doses of cevimeline maintains physiological saliva secretion, acinar cell architecture, and functional innervation long-term

Given that a single dose of ALG+CV at 14 days post-IR was able to improve secretory function, albeit transiently, and reduce the loss of acini and their cholinergic nerve supply, we next asked whether multiple doses could lead to a sustained improvement in glandular function and structure. Due to subcutaneous alginate not degrading over time, with the biogel remaining present at 70 days post-IR, we proceeded to administer the initial treatment of ALG+CV, ALG, or saline at 14 days post-IR, and follow this with weekly local injections of CV into the ALG+CV mice (ALG+multiCV), and saline into the saline or ALG treated mice, starting at 38 days post-IR (i.e., 3 weeks after the initial saline, ALG, ALG+CV at 14 days). This time point was chosen based on saline-treated mice showing a significant reduction in physiological saliva secretion at 41 days post-IR (**Fig.3B**). Physiological salivary function was then measured every 2 weeks from day 35 to 91 post-IR via gustatory stimulation (**Fig.4A**). As for the mice receiving a single dose, both saline-and ALG-treated mice undergoing multiple dosing showed reduced physiological saliva secretion, with a reduction of 62% (*p*<0.001) and 34% (*p*<0.08), respectively, by 49 days post-IR (**Fig.4B**). In contrast, our multidose regimen resulted in ALG+multiCV treated mice having saliva secretion levels that mimicked those of non-IR mice, and were significantly above those of IR mice, for the entire 3-months (91 days) measured (**Fig.4B**), indicating that ALG+multiCV was sufficient to restore physiological saliva secretion long-term.

**Fig. 4.**
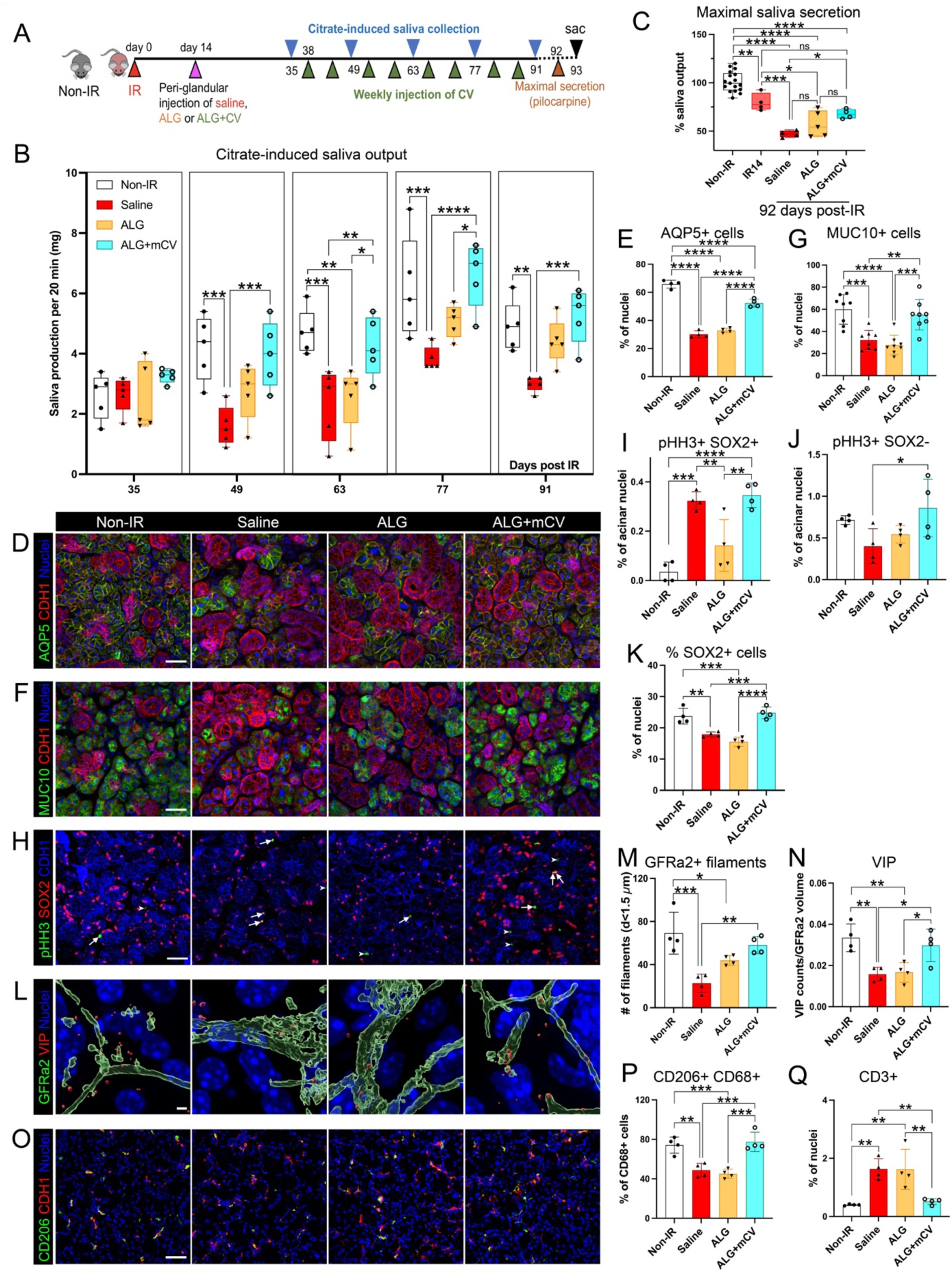
Multiple doses of cevimeline maintains physiological saliva secretion, acinar cell architecture and functional innervation long-term. **A**, Timeline for radiation, treatments and saliva collections. **B**, Citrate-induced saliva output was measured once every 2 weeks from 35 to 91 days post-radiation. **C**, Maximal saliva secretion in response to systemic pilocarpine administration at 92 days. Data was normalized to non-IR controls. **D-G**, Representative images and quantification of AQP5^+^ (**D, E**), and MUC10^+^ (**F, G**) acinar cells. Epithelial cells and nuclei were stained for E-cadherin (CDH1, red) and nuclei (Hoechst 33342, blue). **H**-**K**, Representative images (**H**) and quantification of mitotic (pHH3^+^) acinar progenitor cells (**I**, SOX2^+^) and transit amplifying cells (**J**, SOX2^-^). Quantification of SOX2^+^ acinar cells is shown in **K. L**-**N**, Representative images (**L**) and quantification of parasympathetic nerves marked by GFRa2 (**M**, green, 3D-surface reconstructed with Imaris), and the neuropeptide vasoactive intestinal peptide (**N**, VIP, red, 3D-surface reconstructed using Imaris). Hoechst 33342 = nuclei (blue). GFRa2^+^ filaments with diameters of less than 1.5 µm (innervate the acini) were quantified. VIP was normalized to GFRa2^+^ filament volumes. **O-Q**, Representative images (**O**) and quantification of pro-reparative (CD68^+^CD206^+^) M2-polarizing macrophages (**P**) and CD3^+^ T cells (**Q**). Scale bars in C, E, I and P = 40 µm, in M = 10 µm. Dots in the bar graphs represent biological replicates. Data are expressed as mean ± s.d. \**p* < 0.05; \*\**p* < 0.01, \*\*\**p* < 0.001; a one-way analysis of variance with Dunnett’s multiple comparison test for all graphs except B; a two-way analysis of variance with Dunnett’s multiple comparison test was applied to B.

We next determined the extent to which the acinar compartment was damaged in the different treatment groups by performing a maximal saliva secretion assay through systemic injection of pilocarpine. Both saline- and ALG-treated mice exhibited a significant reduction in maximal saliva production upon stimulation at 92 days compared to non-IR mice (50%;p<0.0001 and 43%;p<0.0001 of non-IR levels, respectively) as well as compared to 14-days post-IR mice (40% reduction (p<0.01) for saline and 29% (p<0.05) for ALG; **Fig.4C**). Although saliva secretion for ALG+multiCV treated mice did not completely mimic those of non-IR controls (30% decrease; p<0.0001), maximal secretory function had a significant 36% increase compared to saline-treated IR mice (*P*<0.05; **Fig.4C**). Furthermore, the maximal secretion level was not significantly different to that of mice at 14-days post-IR (**Fig.4C**), suggesting that ALG+multiCV treatment halts additional gland injury after this time point. Thus, ALG+multiCV substantially aids in the retention of physiological cell function, and is able to maintain maximal secretion from radiation-injured tissue from the time of treatment.

Based on the preservation of saliva output with ALG+multiCV treatment, we next determined whether acinar cell architecture was also positively impacted through immunofluorescent analysis (**Fig.S4A**). As with the day 70-day post-IR submandibular gland, we found the acinar cell compartment to be significantly preserved by ALG+multiCV treatment, with only a 20% reduction in the AQP5^+^ acinar cells compared to non-IR controls (*p*<0.0001), whereas saline and ALG treated tissue showed a 46% and 54% decrease (*p*<0.0001) in acinar cell number, respectively (**Fig.4D, E**). Furthermore, in contrast to the large reduction (>40% decrease) in MUC10-synthesizing acinar cells in saline-(*p*<0.01) and ALG-treated (*p*<0.001) glands, the number of MUC10^+^ cells in ALG+multiCV treated tissue was significantly increased compared to saline-treated IR mice and mirrored that of the non-IR controls (71%;p<0.01; **Fig.4F, G**), suggesting ALG+multiCV serves to maintain secretory protein secreting acinar cells.

Given the extensive preservation of acinar cells with ALG+multiCV treatment (**Fig.4D-G**), we next questioned the impact of the various treatments on the expansion of the global acinar cell pool and the proliferation status of its progenitors at the 93-day post-IR time point. We found that the total number of SOX2^+^ acinar progenitors in saline-(*p*<0.01) and ALG-treated (*p*<0.001) IR tissue was significantly lower than non-IR control glands (**Fig.4K**), confirming that acinar progenitors are not maintained post-IR. Unexpectedly, we measured a significant increase in proliferating pHH3^+^SOX2^+^ cells in saline-treated IR glands as compared to non-IR tissue (10-fold,p<0.001, **Fig.4H**,**I**). However, the number of pHH3^+^SOX2^-^ transit amplifying cells was reduced (**Fig.4J**), suggesting that SOX2^+^ progenitors do not undergo active cell division to produce differentiated daughter cells. In contrast to saline or ALG treatment, ALG+multiCV-treatment not only significantly increased the number of pHH3^+^SOX2^+^ progenitors (12-fold compared to non-IR glands; p<0.0001; **Fig.4H,I**), but also expanded the number of pHH3^+^SOX2^-^ transit amplifying cells (2-fold increase over saline-treated IR glands, p<0.05, **Fig.4J**) while maintaining the SOX2^+^ progenitor population equal to non-IR glands. Thus, these data indicate that ALG+multiCV is not only able to increase the pool of actively dividing SOX2^+^ progenitors for the repopulation of the progenitor cell pool, but also ensures a large population of proliferating transit amplifying cells that ultimately replenishes the acinar cell compartment.

Similar to our analysis at 70 days post-IR, we found ALG+multiCV-treatment to substantially rescue functional parasympathetic innervation of secretory acini. Indeed, innervation of acinar cells by thin GFRa2^+^ filaments (diameter < 1.5μm) closely resembled that of non-IR glands (**Fig.4L and Fig.S4C**) whereas GFRa2^+^ filaments in saline-treated tissue only reached 40% of non-IR levels (*p*<0.001; **Fig.4M**). Furthermore, consistent with the gustatory-stimulated saliva secretion outcomes (**Fig.4B**), parasympathetic nerve function was also maintained, as shown by the levels of vasoactive intestinal peptide (VIP), a key pro-secretory neuropeptide (*47*), being similar in both ALG+multiCV-treated and non-IR salivary glands, and significantly greater than the saline IR controls (2-fold; p<0.01; **Fig.4L, N and Fig.S4D**). Although ALG treatment also showed some increase in innervation of acini (**Fig.4L, M and Fig.S4D**), the levels of VIP resembled that of saline-controls (**Fig.4N and Fig.S4D**), indicating ALG alone does not rescue nerve function.

Finally, we questioned immune cell infiltration in response to the different treatments at 93 days post-IR. Similar to the 70-day post-IR tissue, IR significantly increased the total CD68^+^ pan-macrophage population within the glands under all treatment conditions (*P*<0.001 for saline and ALG+multiCV; p<0.0001 for ALG; **Fig.S4B**). However, only ALG+multiCV treated salivary glands contained significantly higher numbers of pro-reparatory CD206^+^ M2 macrophages (**Fig.S4B**), with the percentage of CD206^+^CD68^+^ macrophages (of total CD68^+^ cells), being significantly increased compared to saline-IR salivary glands (60% increase; p<0.001) and similar to non-IR controls (**Fig.4O, P**). We also found the number of CD3^+^ T cells in ALG+multiCV treated tissue to resemble that of non-IR controls, contrasting greatly with the substantial infiltration of CD3^+^ T cells (3-fold increase; p<0.01) into saline-and ALG-treated glands (**Fig.4Q and Fig.S4E**).

Taken together, these studies indicate that extending the delivery time of cevimeline is sufficient to maintain long-term physiological saliva output, improve tissue architecture and promote a pro-repair environment in response to radiation damage (**Fig.5**).

**Fig. 5.**
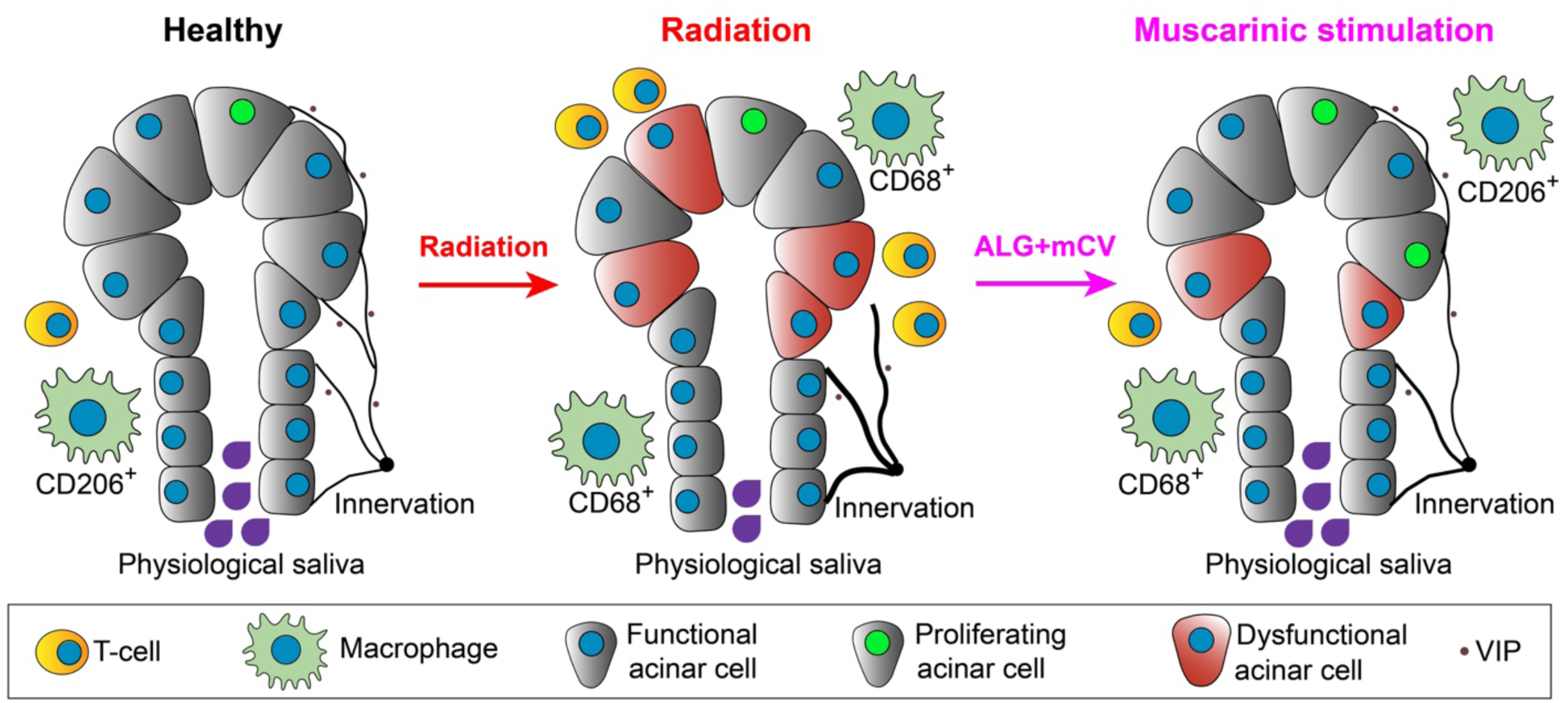
Model of muscarinic-induced acinar cell regeneration. Under healthy conditions, salivary acinar cells (green) are highly secretory and extensively innervated by parasympathetic axons (black) that regulate both cellular function and tissue homeostasis through the production of neurotransmitters e.g., VIP (black dots). Upon radiation damage, salivary gland acinar cells undergo degeneration, acinar progenitor cells are reduced in number, functional parasympathetic innervation of acini is lost, and the tissue is infiltrated by T cells and macrophages that are polarized toward pro-inflammatory phenotypes (e.g., CD68^+^CD206^-^). However, treatment of irradiated glands with the muscarinic agonist, cevimeline, encapsulated in an alginate hydrogel reverses these outcomes, resulting in acinar cell regeneration, the maintenance of acinar progenitor cells, a pro-repair immune response and the restoration of innervation and glandular function.

## Discussion

Despite the vast array of tissues inadvertently damaged by radiation treatment for cancers, with salivary glands being one of the most frequently injured tissues (*48*), regenerative therapies capable of restoring organ architecture and physiological function have remained elusive. We, and others, have demonstrated that nerves are critical regulators of somatic stem cell behavior (*10, 49, 50*) but - to date - it was unknown whether they also mediate regenerative responses after radiation-induced damage and if so, by what mechanism(s). Here, we show that local administration of a muscarinic receptor agonist to irradiated salivary glands promotes acinar cell replacement, maintains tissue architecture and parasympathetic innervation, and restores secretory function over an extensive time course, thereby preserving physiological levels of saliva necessary for oral and systemic well-being (see **Fig.5**). Together, our data indicate that irradiated epithelial tissues have the capacity to respond to pro-regenerative treatments, potentially months after radiation treatment in humans, and support the application of this novel injectable hydrogel therapy for reversing salivary gland dysfunction after radiation induced damage.

Acinar cell degeneration is the most prominent outcome of radiation-induced damage to salivary glands (*6, 40*), resulting in the loss of saliva protein synthesis and fluid secretion. Surprisingly, however, very few studies to date have been actively engaged in directly promoting acinar cell replacement after radiation damage. Rather, strategies have been based on preserving acinar cells through inhibition of apoptosis (*51*) and senescence (*52*), restoring healthy tissue through the application of stem cell-derived organoids that contain acinar cells (*53, 54*), and the deletion of damaged cells (*55*). Here, we demonstrate that the acinar cell pool can be directly regenerated through activation of muscarinic receptors enriched on the acinar progenitors, thus repopulating the glands with functional secretory acini. Muscarinic receptor signaling has previously been shown to promote stem cell expansion (*10, 56*) and epithelial cell division (*56, 57*) in other organ systems. Specifically in the salivary gland it induces extensive acinar cell proliferation in un-injured conditions *in vivo* (*10*), as well as in *ex vivo* explants immediately after radiation (*10*). However, this is the first *in vivo* study to reveal that acinar cell replacement can be activated weeks to months after radiation damage, preventing further degeneration of the acinar compartment and preserving physiological saliva secretion.

Little is known about the ability of acinar cells to renew after radiation, other than their capacity to undergo increased cycling up to 20 days post-IR (*58*). This cellular response has been confirmed by genetic lineage tracing studies in mice where recombination is activated in acinar cells before IR, with glands showing active replenishment of acinar cells within 14-days (marked by *Sox2* (*10*)) or up to 90 days (marked by *Mist1* (*59*)) post-IR. Human salivary gland acinar cells may also have regenerative potential, as, despite the severe loss in acini at 2.4 months post-IR, small clusters of acinar cells still express proliferative and functional markers after >10 years post-radiotherapy treatment (*8*). However, in all cases, the endogenous replenishment is not sufficient to maintain organ structure and function post-IR, with all rodents and human patients ultimately showing loss of saliva output. Our findings confirm that such spontaneous proliferation in SOX2^+^ acinar cells can occur up to 70-90 days post-IR, but is unable to maintain the overall SOX2^+^ acinar progenitor cell pool necessary to repopulate the injured gland. To date, it remains unclear what signal(s) induce the initial cycling of acinar progenitors after radiation, and why this is lost over time. Our data demonstrating that the maintenance of muscarinic receptor activation in irradiated glands is sufficient to sustain acinar cell proliferation and the overall acinar cell pool over the long term, and the known pro-proliferative role of muscarinic agonism, suggests this pathway is a key regulator of cell regeneration. However, further studies are needed to determine why endogenous muscarinic agonism fails as well as if other signaling systems are involved. Furthermore, whether removal of cevimeline after 3 months of treatment returns the gland to a self-sustaining state requires further investigation.

Not only is replenishment of the acinar cell pool essential for tissue repair, it also requires maintenance of the surrounding nerve-enriched niche that influences secretion and organ homeostasis (*41*). With the ablation of parasympathetic nerve function resulting in acinar cell atrophy and the absence of saliva (*10, 60, 61*), the severe reduction in parasympathetic innervation of human salivary glands 2 years after radiation therapy (*62*), and the reduction in acinar cell innervation 2-3 months after radiation, as revealed in this study, is likely a significant mediator of irreversible degeneration. The ability of CV, when administered weekly, to preserve this essential, functional interaction allows for the acquisition of a near homeostatic state. How CV acts to sustain this positive feedback loop between nerves and acini remains unclear. It is well known that nerve-target cell interactions are initiated and maintained by neurotrophic factors produced by the target cells (*63*), including acinar cells. The neurotrophic factor neurturin (NRTN), is key a regulator of acinar cell innervation by parasympathetic nerves during salivary gland development (*62*), it increases innervation when overexpressed in irradiated adult tissue (*64*), and is significantly reduced in irradiated adult human acinar cells (along with the nerves) (*62*). Although it is not known whether muscarinic receptors specifically regulate NRTN production, muscarinic agonists have been shown to modulate the production of other neurotrophins, such as NGF, in gastric epithelial (*65*), adipose (*66*) cells to promote innervation, suggesting this as a potential mechanism by which cevimeline acts in the salivary glands. Whether this is the case remains to be determined.

Given a chronic pro-inflammatory response to radiation damage is a major contributor to human salivary gland dysfunction and degeneration, with irradiated human salivary glands exhibiting extensive lymphocytic infiltration (*67*), controlling this outcome is also required to enable robust tissue regeneration. We found that cevimeline treatment reduced the pro-inflammatory state and instead, promoted a pro-repair immune response in the salivary glands after radiation, an outcome consistent with a regenerative rather than degenerative response. The cholinergic system has previously been described as anti-inflammatory, with a growing number of studies reporting neuronal acetylcholine to attenuate the production of pro-inflammatory cytokines via α7 nicotinic receptors on macrophages, microglia and astrocytes (*68*). However, the role of muscarinic receptors in the regulation of immune cell identity and function remains poorly understood. Although a number of previous studies primarily link muscarinic signaling in immune cells to pro-inflammatory outcomes, including the expansion of macrophages (*69*), more recent studies have suggested an anti-inflammatory role. In the murine small intestine, genetic deletion of *Chrm3* was associated with an upregulation of pro-inflammatory T_H_1/T_H_17 cytokines in the absence of infection (*70*). Furthermore, upon infection with the nematode, *Nippostrongylus brasiliensis, Chrm3*^*-/-*^ mice were unable to mount the appropriate anti-inflammatory T_H_2 responses required to efficiently clear the infection (*70*). This is significant because M1 (pro-inflammatory) macrophages promote T_H_1 responses, which then suppress M2 (pro-reparatory) macrophage polarization, and vice versa, with M2 macrophages and T_H_2 responses suppressing M1 macrophage polarization (*71*). Consistent with this outcome, we show that CV treatment, as opposed to ALG or saline, results in the specific expansion of the M2 subtype as well as reduces T cell infiltration, thus supporting muscarinic agonism as a mitigator of inflammation and a promoter of wound healing under radiation conditions. However, further studies are required to confirm that this is indeed the case.

Taken together, our muscarinic based alginate treatment is able to produce a functional secretory tissue with preserved innervation after radiation exposure. The beneficial cellular changes that we observed at both the acinar cell and nerve levels, and importantly, the maintenance of saliva secretion itself, demonstrates that this is a promising treatment for the functional improvement in saliva output in those suffering from radiation-induced xerostomia.

## Materials & Methods

### Animal studies

Adult female mice (6-7 weeks) from the C57BL/6 inbred strain (Harlan Laboratories) were used in all experiments. Mice were socially housed and monitored for 48 h following radiation procedure. Animals received either no radiation or a single 10 Gy dose, as described below. At the end of experiments, animals were euthanized using CO_2_ at a flow of 2 L/min followed by a cervical dislocation.

### γ-Radiation of salivary glands

Mice were treated with a single 10 Gy dose of γ-radiation 14 days before any treatment was given, as performed previously (*10, 72*). Animals were anaesthetized with 2.5 % Avertin (2,2,2-Tribromoethanol (Alfa Aesar), tert-Amyl Alcohol (2-Methyl-2-butanol, Spectrum) in 0.9 % saline (Vedco Inc, 100µl/10g body weight.). Animals were placed into a Shepherd Mark I Irradiator (JL Shepherd & Associates), with the body and cranium shielded from radiation using lead blocks. The head and neck regions of mice were exposed to two doses of 5 Gy at a dose rate of 167 Rads/min for 2.59 min (one of each side of the head, bilateral, and sequential but on the same day) for a total dose of 10 Gy, to irradiate the salivary glands. This dose was calculated by isodose plot mapping (dose distribution), provided by the manufacturer. Control mice were anesthetized as per experimental mice but did not undergo radiation treatment. All mice were allowed to completely recover before returning to normal housing and were given soft diet ad libitum (Clear H_2_O).

### Peri-glandular application of alginate+cevimeline or alginate-only 14 days after radiation exposure

*In vivo* delivery of alginate (control group) versus alginate+cevimeline (treatment group) was achieved via subcutaneous injection superficial to both the sublingual and submandibular salivary glands without surgical exposure (as detailed above) 14 days after radiation-induced damage. Oxidized alginate hydrogels (2 % at 5 wt%) were prepared as described previously (*73*). Immediately before *in vivo* delivery lyophilized alginate (50 mg, gift from Eben Alsberg) was dissolved in DMEM (894.2 µL; Gibco #11885084) containing cevimeline (84.8 µl of 100 mM cevimeline, Sigma-Aldrich, SML0007) or sterile RNase free water (84.8 µL; Invitrogen #10977015) and crosslinked with 100 µL of supersaturated CaSO4 (84 mg/ml). Animals were anesthetized using isoflurane (2.5 % induction, 1.5 % maintenance), skin was cleaned with alternating iodine and alcohol washes, and 50 µL of the alginate alone, alginate+cevimeline or saline were administered by subcutaneous injection using a 25G x 5/8 needle placed at an angle of 45° to the tissue. The location of each injection was 0.5 cm to the left or to the right of the midline, between c2 and c5 vertebra. To ensure correct localization of the alginate hydrogels around the salivary gland, Iohexol (Milipore Sigma 74147), a radio-opaque contrast reagent, was encapsulated into the hydrogels at a concentration of 0.35 g/mL and a Hologic Fluoroscan Premier Encore 60000 C-Arm Imaging System (45 kV, 0.031 mA) was used to visualize the material. Mice were euthanized at the end of experiments (n=3-6) and the sublingual and submandibular glands processed for immunofluorescent analysis, as described below.

### Physiological saliva collection (gustatory-stimulated)

Physiological saliva secretion was measured using gustatory stimulation, as previously published with some minor modifications (*31*). Briefly, 7.5 × 10 cm filter paper (Bio-Rad, 1703932) was incubated in 73.6 mg/ml sodium citrate (Spectrum Chemical, S1250) solution for 15 min at room temperature (RT) on an orbital shaker and then dried overnight at RT. The filter paper was then cut into 30 mm x 2 mm strips and the middle point labelled on each strip. Individual strips were then placed in pre-weighed 1.5ml microcentrifuge tubes. After mice were anesthetized by 2 % inhaled isoflurane, a strip of filter paper was inserted into the oral cavity on top of the tongue until the maxillary incisors reached the middle point of the strip. After 20 min the strip was removed and placed in the original 1.5 ml microcentrifuge tube. The amount of saliva collected was determined by measuring the difference in weight of the microcentrifuge tube with filter paper before and after collection, using a precision scale (OHAUS Adventurer).

### Maximum saliva collection (pilocarpine-stimulated)

Maximum saliva output was measured as described previously (*74*). Briefly, 68ng/µl pilocarpine (Sigma-Aldrich, P0472) in saline (Aspen, AHI14208186) solution was freshly prepared before saliva collection. Mice were anesthetized by 2 % inhaled isoflurane, and pilocarpine (200 µl/30 g body weight of 68 ng/µl) injected via i.p., then animals were placed back in the anesthetizing chamber for 3 minutes. Pre-weighed cotton balls (Puritan, 806-WCL) were inserted into the mouths of the mice. After 10 minutes, one cotton ball was exchanged for another, and left for another 10 minutes; in total, saliva was collected for 20 minutes using 2 cotton balls. These balls were placed back in the original 1.5 ml microcentrifuge tube and the amount of saliva collected was determined by measuring the difference in weight of the cotton balls before and after collection, using a precision scale (OHAUS Adventurer).

### Preparation of tissue samples for RNAseq and immunofluorescent analyses

For RNA analysis, tissue was snap frozen and stored at −80 °C. For immunofluorescent analysis, tissue was either freshly frozen in optimal cutting temperature compound (OCT) (Tissue-Tek) and stored at −80 °C, or immediately fixed with 4% paraformaldehyde (PFA) overnight at 4°C. Fixed samples were washed with PBS, cryoprotected by immersion in 12.5% and 25% sucrose solution, and then embedded in OCT and stored at −80 °C. Tissues were sectioned (12µm) with a cryostat (ThermoFisher Scientific) for immunofluorescence staining described below.

### Bulk RNA sequencing of salivary glands

RNA was isolated from 4 × 50 µm tissue sections of non-IR, 7- or 14-days post-IR sublingual glands using the RNAqueous Micro Kit (ThermoFisher Scientific, AM1931) and total RNAs were treated with DNase I (ThermoFisher Scientific). RNA integrity and quantification were assessed using the RNA Nano 6000 Kit of the Bioanalyzer 2100 system (Agilent Technologies, CA). RNA sequencing was performed by Novogene (https://en.novogene.com) using Illumina NovaSeq 6000 platform. Raw data (FASTQ) were processed through fastp. Paired-end clean reads were aligned to the reference genome using the Spliced Transcripts Alignment to a Reference (STAR) software (*75*). FeatureCounts was used to count the read numbers mapped to each gene (*76*). Differential expression analysis between two conditions was performed using DESeq2 R package (*16*). Genes with an adjusted p-value < 0.05 were assigned as differentially expressed. Gene Ontology of differentially expressed genes was analyzed using g:Profiler (*17*).

### EdU (5-ethynyl-2’-deoxyuridine) delivery and detection

To quantify the rate of cell turnover, non-IR and IR animals were systemically injected with 0.9% saline containing 0.25 mg per 25 g of body weight of 5-ethynyl-2’-deoxyuridine (EdU, Thermofisher Scientific) followed by a 2 hour or 7 days chase before sacrifice. EdU detection was performed using the Click-iT^®^ EdU Imaging Kit (ThermoFisherScientific, C10086) according to manufacturer’s instruction. Briefly, fresh-frozen sections were fixed with 4% PFA for 15 min at RT followed by two washes with 3% BSA in PBS. Immediately after BSA washes, the sections were immersed in freshly prepared reaction cocktail (430µl 1x Click-iT^®^ reaction buffer + 20µl CuSO_4_ + 1.2 µl Alexa Fluor^®^ azide + 50µl Reaction buffer additive, in proportion). After 30 minutes’ incubation, sections were washed twice with 3% BSA in PBS. Images were obtained with an LSM900 confocal microscope.

### Immunofluorescence studies

Tissue section immunofluorescence analysis was completed as described previously (16). In brief, fresh-frozen tissue was fixed with 4% paraformaldehyde (PFA) for 10 min, or pre-fixed tissue were defrosted for 5 min, followed by permeabilizing with 0.5% Triton-X100 in PSB for 10min at RT. Tissue sections were blocked for 2 hours at RT with 10% Donkey Serum (Jackson Laboratories, ME), 5% BSA (Sigma Aldrich), and MOM IgG-blocking reagent if required (Vector Laboratories, CA) in 0.01% PBS-Tween-20. Salivary gland sections were incubated with the following primary antibodies overnight at 4 ºC: stem cell antibody goat anti-SOX2 (1:200, Neuromics, GT15098); epithelial cell antibody, rat anti-E-Cadherin (1:400, Invitrogen, 13-1900); cell proliferation antibody, rabbit anti-pHH3 (1:200, Cell Signaling Technology, #9701); muscarinic M3 receptor (CHRM3) antibody, rabbit anti-CHRM3 (1:500, Research Diognostics, AS-3741S); acinar antibodies rabbit anti-AQP5 (1:200, Millipore, AB3559), anti-MUC10 (1:200, Abcore, AC21-2394); immune cell antibodies, rat anti-CD68 (1:400, Biorad, MCA1957), rabbit anti-CD3 (1:200, AbCam, ab5690), and rabbit anti-CD206/MMR (1:200; AF2535; R&D systems); parasympathetic nerve/ Schwann cell marker, goat anti-GFRα2 (1:200; R&D Systems AF429); and vasoactive intestinal peptide antibody anti-VIP (1:100, Abcam, ab124788). Antibodies were detected using Cy2-, Cy3-or Cy5-conjugated secondary Fab fragment antibodies (1:300, Jackson Laboratories) and nuclei stained using Hoescht 33342 (1:1000, AnaSpec Inc.). Slides were mounted using Fluoromount-G (SouthernBiotech) and images were taken using an LSM900 confocal microscope. Image processing and quantification was performed using NIH ImageJ software or Imaris v9.6 (Bitplane). For image analysis, 3-5 fields of 250 µmx 250 µm sections (1 µm z sections) were imaged within different sections of the tissue.

### Quantification of immunofluorescent images and specific cell populations

Acinar cells were identified based on NKCC1 or AQP5 expression, proliferating cells were identified based on EdU incorporation or pHH3 expression and immune cells were identified based on the T cell marker CD3 or the macrophage markers CD68 and CD206. Quantification of cells was achieved by applying the image processing software ImageJ (NIH) and manually counting cells. Acinar cell boundaries were determined by AQP5, NKCC1 or Ecadherin (CDH1) expression. The number of acinar cells was normalized to the total number of nuclei per section, and the number of proliferating acinar cells was normalized to the total acinar cell number per region. The number of immune cells was normalized to the total number of nuclei per section. Each quantification was performed on 3-4 sections per gland, from at least 3 animals.

Quantification of GFRa2^+^ parasympathetic nerves and the neuropeptide VIP was achieved through analysis of 256 µm x 256 µm images within a 12 × 1 µm stack. Three separate regions of each gland that were enriched in acinar cells were imaged per animal (n = 3). Using Imaris v9.6 (Bitplane), images were subjected to Gaussian filtering and background subtraction. Surface reconstructions were made with the “Surfaces” module. GFRa2^+^ nerve fibers were manually traced with the “Filaments” module, “AutoPath” function. Mean volumes and filament diameters were exported from the statistics tab and analyzed in Microsoft Excel and GraphPad Prism. VIP was measured with the “Spots” module, with a distance less than 4.5µm to the GFRa2^+^ nerve fibers. Total GFRa2^+^ nerve fibers and VIP counts were exported from the statistics tab and analyzed in Microsoft Excel and GraphPad Prism.

### Statistical analyses

Statistical tests were performed using GraphPad Prism software v8. Data are plotted as individual data points with mean ± standard deviation. Groups with only two datasets were analyzed with a 2-tailed unpaired Student’s t test. For multiple comparisons an ordinary one-way ANOVA was used to determine if significant differences existed, followed by either a Tukey’s Hardly Significant Difference (HSD) post-hoc comparisons test (for comparing means of multiple groups) or a Dunnett’s test. Significance was assessed using p-value cutoffs indicated as follows: *p < 0.05, **p < 0.01, ***p < 0.001, and ****p < 0.0001. Specific data set analyses are described in the figure legends.

### Study approval

All procedures were approved by the UCSF Institutional Animal Care and Use Committee (IACUC) and were adherent to the NIH Guide for the Care and Use of Laboratory Animals.

## Acknowledgements

The authors would like to thank Drs. Licia Selleri, Jeffrey Bush and Ophir Klein for their contributions to the manuscript.

## Funding

National Institute of Dental and Craniofacial Research (NIDCR; 1R35DE028255) NIDCR C-DOCTOR (U24DE026914)

Tobacco Related Disease Research Program (TRDRP; 588359).

## Author Contributions

Conceptualization: IMAL, CSB and SMK

Methodology: JL, SS, LB, AM, SM, EG, NCP and EA

Investigation: JL, SS, LB, AM, SM, EG and NCP

Formal analysis: JL, SS, LB, AM, SM, EG and NCP

Visualization: JL, SS, LB and SMK

Data curation: JL, SS and LB

Providing reagents: EA, OJ

Project administration: SMK

Funding acquisition: SMK

Supervision: SMK

Writing- original draft: JL, SS, LB, IMAL, CSB and SMK

Writing- review & editing: JL, SS, LB, IMAL, CSB and SMK

## Competing interests

The authors declare that they have no competing interests.

## Supplementary Materials (Separate file)

Please see the attached “Li, Sudiwala et al 2022 Supplementary Figures.pdf” for supplementary materials.

## Notes

### Competing Interest Statement

The authors have declared no competing interest.

