## Supplementary Data for "Long-term functional regeneration of radiation-damaged salivary glands through delivery of a neurogenic hydrogel"

**This PDF file includes:**

Figs. S1 to S4
Data S1 to S3

**Fig. S1.**

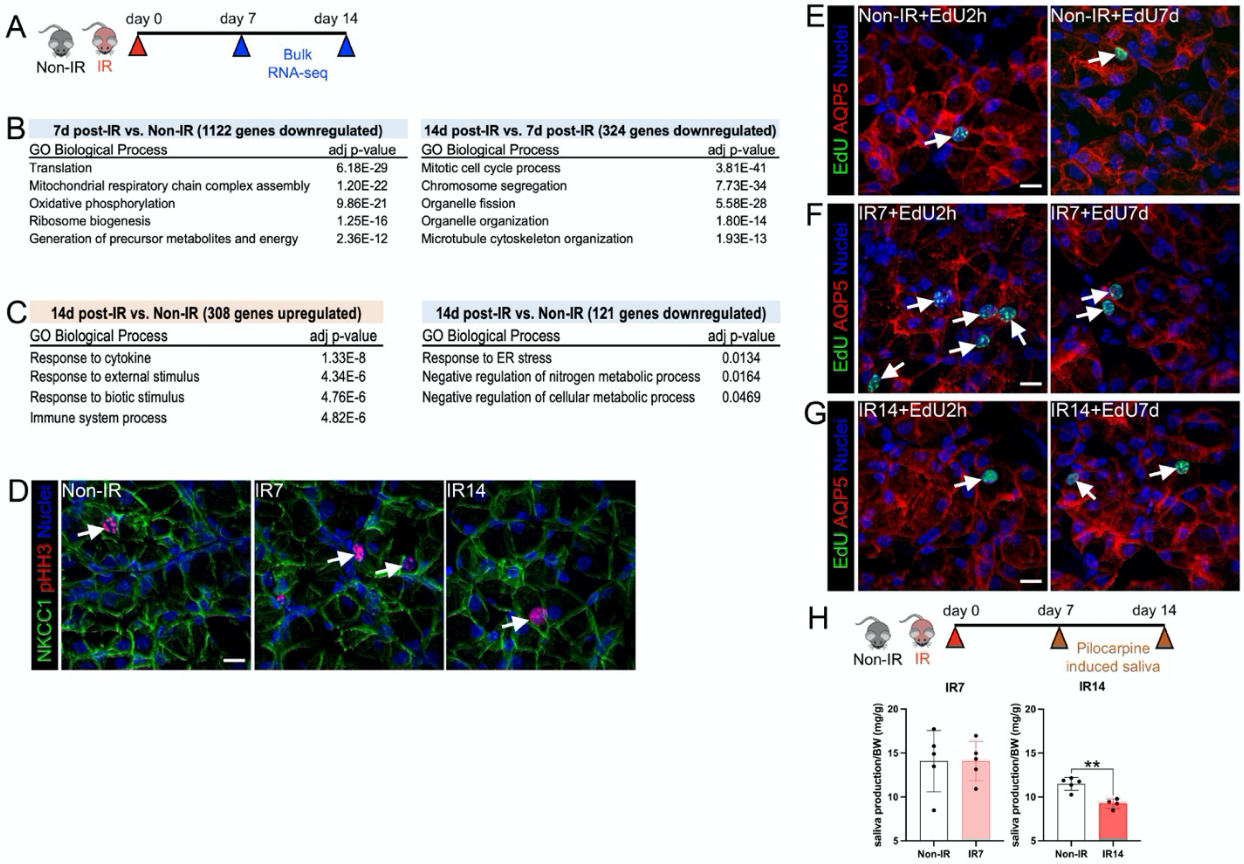


**Fig. S1. Transcriptomic and functional analysis of sublingual glands at 7- and 14-days post-IR. A**, Timeline of radiation exposure and tissue collection for bulk RNA-seq. **B**, Gene ontology (GO) analysis of down-regulated genes within sublingual glands in response to radiation. Transcriptomes from 7-days post-IR (IR7) were compared to non-IR or to 14-days post-radiation (IR14), n=3 for each group. **C**, Gene ontology (GO) analysis of up- and down-regulated genes within sublingual glands 14 days post-IR, compared to Non-IR control, n=3 for each group. **D**, Sublingual glands from non-IR (control), 7- and 14-days post-IR mice were immunostained for the mitotic marker phospho-histone H3 (pHH3, red) and acinar marker, Na-K-Cl cotransporter (NKCC1, green). White Arrows point to pHH3$^{+}$ acinar cells. **E-G**, Immunofluorescent detection of EdU$^{+}$ (green) acinar (AQP5, red) cells in sublingual glands at 2 h and 7 days following a single injection of EdU into non-IR mice (**E**), and mice at 7- (**F**) and 14-days post-IR (**G**). White Arrows point to $EdU^+$ acinar cells. **H**, Maximal saliva output from murine salivary glands was conducted through performing a systemic injection of pilocarpine and saliva was collected 7- or 14 days post-IR. Non-IR mice were used as controls. N=4-5 mice per condition. Each dot in the bar graph represents a biological replicate. Data are expressed as mean ± s.d. An unpaired two tailed t-test was applied to graphs in H, $^{**}P<0.01$. Scale bars= 40 $\mu$m.

**Fig. S2.**

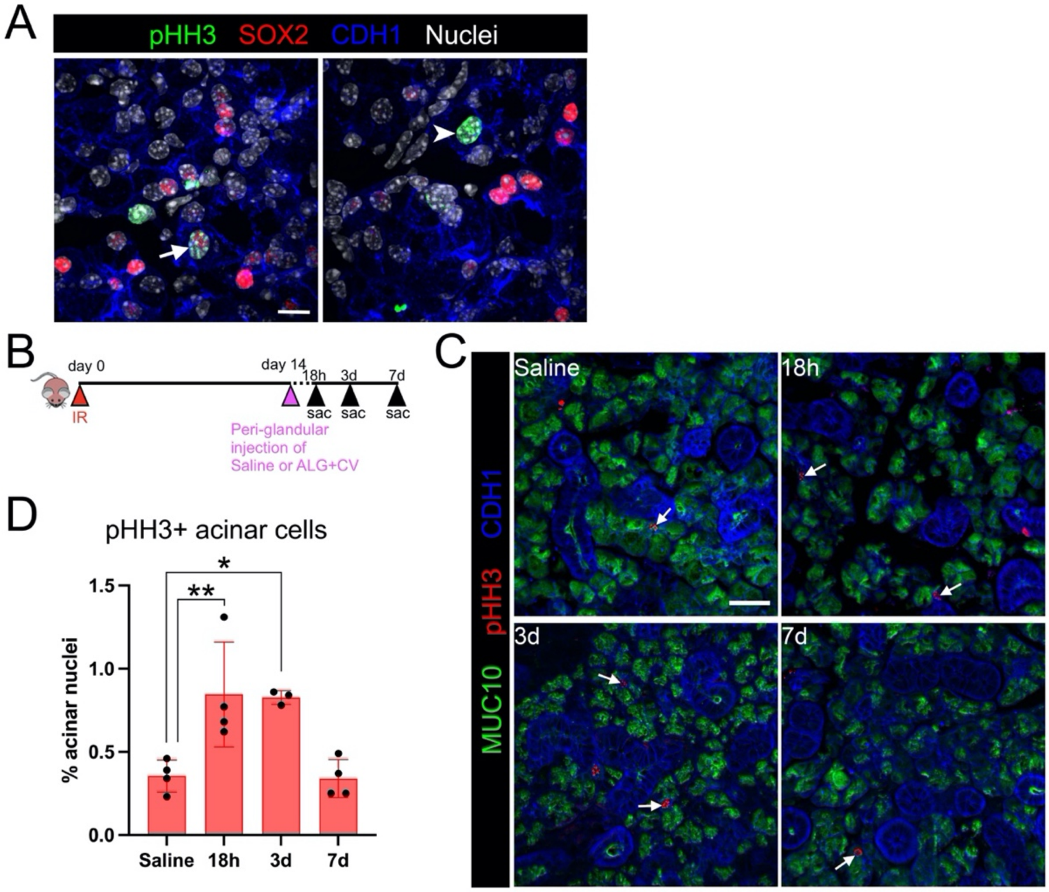


**Fig. S2. Cevimeline incorporated into alginate can promote acinar proliferation in salivary glands after radiation. A,** Immunostaining of mitotic marker phospho-histone H3 (pHH3, green), progenitor marker (SOX2, red) and epithelial cell marker (CDH1, blue) in the sublingual gland. A white arrow points to a pHH3$^{+}$ SOX2$^{+}$ cell, a white arrowhead points to a pHH3$^{+}$ SOX2$^{-}$ cell. **B**, Schematic showing treatment regimen in mice**. C,** Immunofluorescent analysis of the serous acinar cell marker mucin 10 (MUC10, green), mitotic marker (pHH3, red) and epithelial marker E-cadherin (CDH1, blue). Mice were treated with ALG+CV and compared to saline controls. White arrows point to pHH3$^{+}$ acinar cells. **D**, Graph indicates the number of proliferating acinar cells.

Scale bar in A = 10 μm, in C = 40 μm. Dots in the bar graph represent biological replicates. Data are expressed as mean ± s.d. One-way ANOVA with Dunnett's multiple comparison test, *$P$<0.05; **$P$<0.01.

**Fig. S3.**

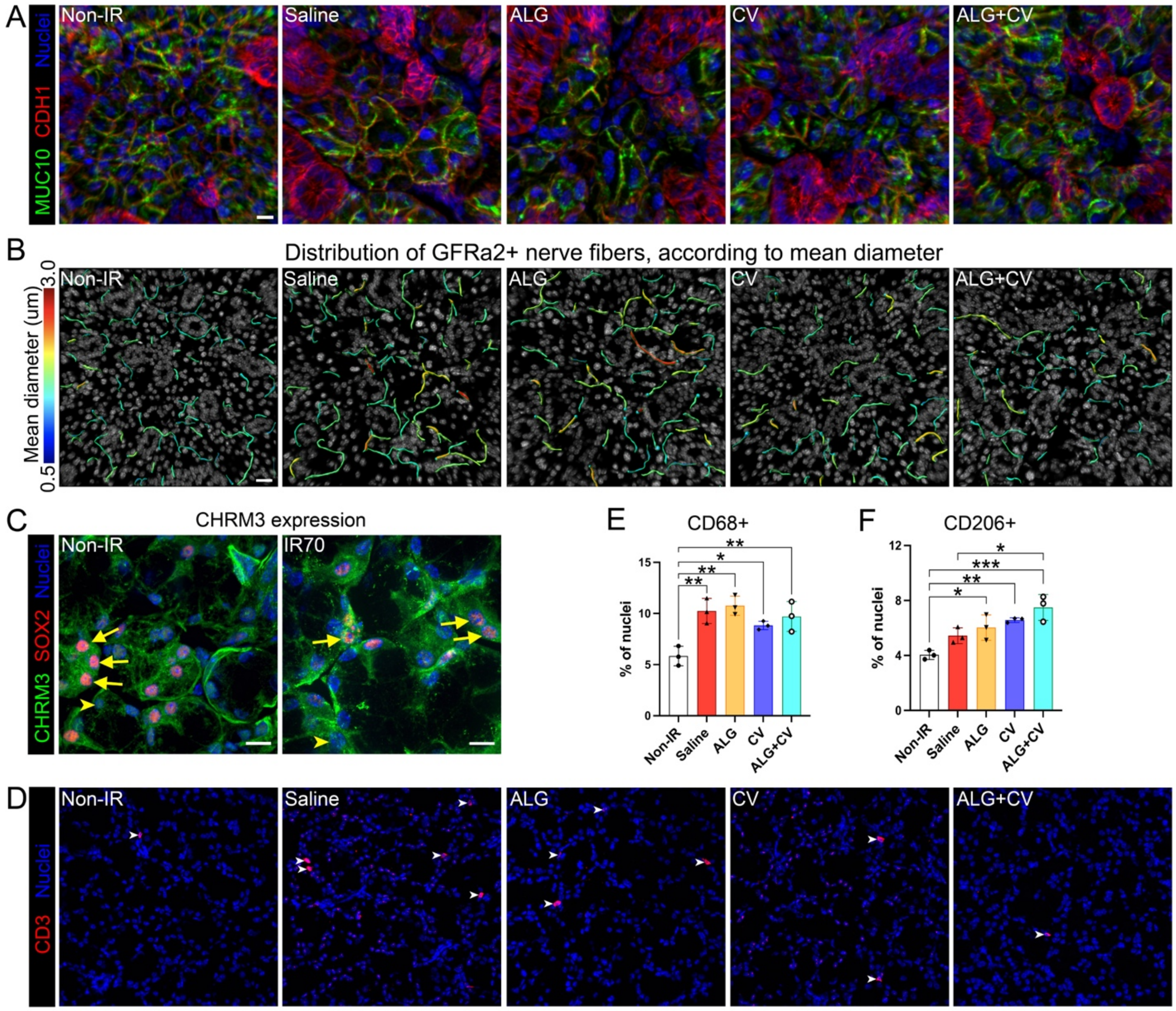


**Supplementary Figure 3. A single-dose of ALG+CV can promote acinar proliferation and ameliorate inflammation after radiation-induced damage. A**, Immunostaining of acinar marker aquaporin 5 (AQP5, green), epithelial marker E-cadherin (CDH1, red) and nuclei (Hoechst 33342, blue). **B**, Mean diameter distribution of GFRa2$^{+}$ nerve fibers in submandibular glands. Nerve fibers were traced with "AutoPath", Imaris. **C**, Representative image of sublingual gland tissue immunostained for CHRM3 (green), SOX2 (red) and nuclei (blue) at 70 days post-IR. Yellow arrows = CHRM3$^{+}$ SOX2$^{+}$ cells. Yellow arrowheads = CHRM3$^{+}$ SOX2$^{-}$ cells. **D**, Representative images of tissue immunostained for $CD3^+$ T cells (red). Nuclei in blue (Hoechst 33342). **D-E**, Quantification of total number of $CD68^+$ (**E**) and M2 $CD206^+$ (**F**) macrophages. Dots represent biological replicates. Scale bars in A and C = 10 μm, in B = 20 μm and D = 40 μm. Data are expressed as mean ± s.d. A one-way ANOVA with Dunnett's multiple comparison test was applied to C and D, $^*p < 0.05$; $^{**}p < 0.01$, $^{***}p < 0.001$.

**Fig. S4.**

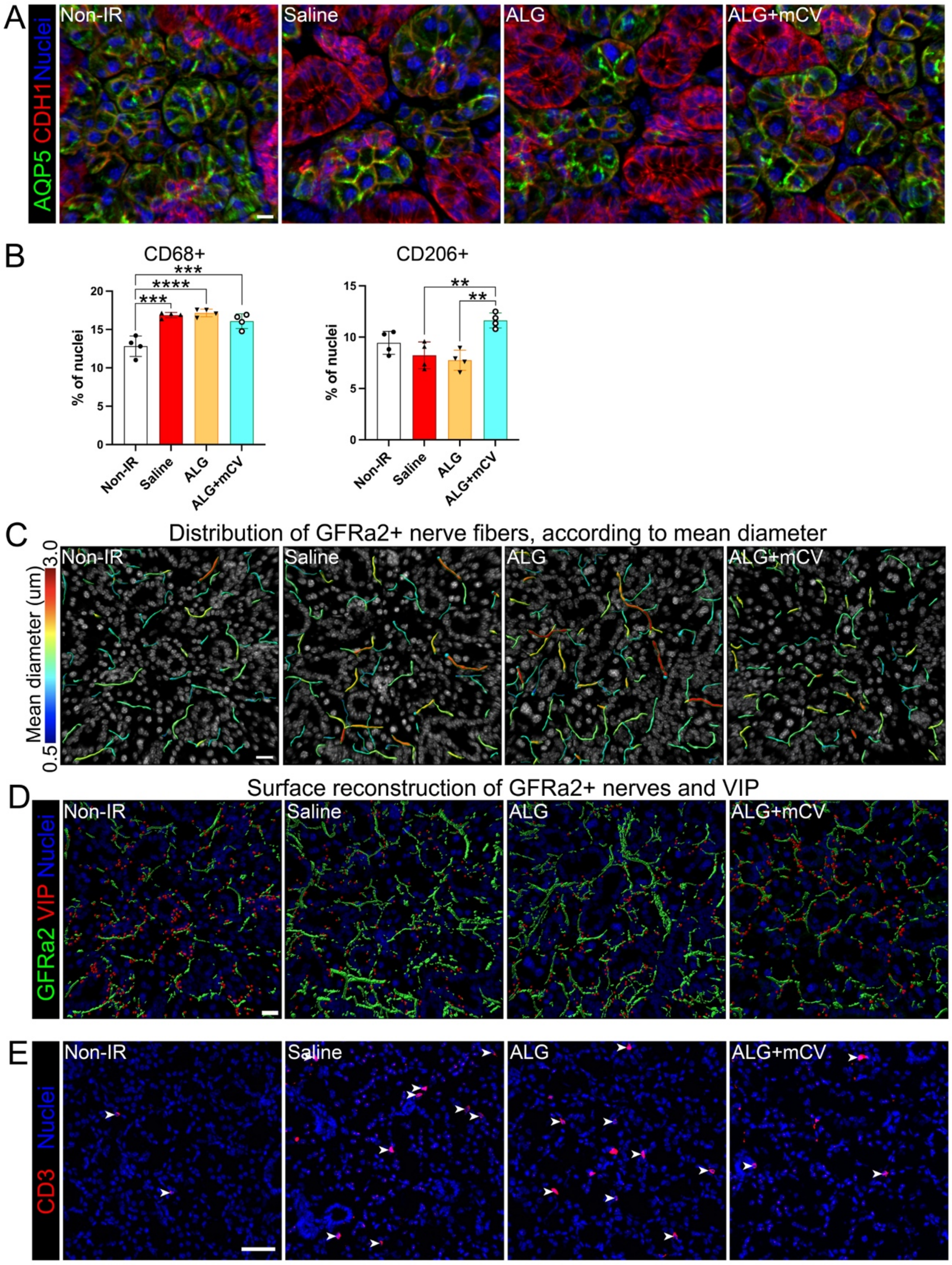

A
AQP5 CDH1 Nuclei
Non-IR
Saline
ALG
ALG+mCV
B
CD68+
% of nuclei
***
****
***
20
15
10
5
0
Non-IR
Saline
ALG
ALG+mCV
CD206+
% of nuclei
**
**
15
10
5
0
Non-IR
Saline
ALG
ALG+mCV
C
Distribution of GFRa2+ nerve fibers, according to mean diameter
Mean diameter (um)
3.0
0.5
Non-IR
Saline
ALG
ALG+mCV
D
Surface reconstruction of GFRa2+ nerves and VIP
GFRa2 VIP Nuclei
Non-IR
Saline
ALG
ALG+mCV
E
CD3 Nuclei
Non-IR
Saline
ALG
ALG+mCV

**Fig. S4. Multiple-dose of ALG+CV ameliorate inflammation after radiation-induced damage in mouse sublingual salivary glands. A**, Immunostaining of acinar cells (AQP5, green), epithelial cells (CDH1, red) and nuclei (Hoechst 33342, blue). **B**, Quantification of total number of macrophages ($CD68^+$) and pro-repair M2 macrophages ($CD206^+$). **C**, Mean diameter distribution of $GFRa2^+$ nerve fibers in submandibular glands. Nerve fibers were traced with "AutoPath", Imaris. **D**, Representative images of parasympathetic nerves marked by GFRa2 (green, 3D-surface reconstructed with Imaris), and the neuropeptide vasoactive intestinal peptide (VIP, red, 3D-surface reconstructed using Imaris). Hoechst 33342 = nuclei (blue). **E**, Immunostaining of T cells (CD3, red) and nuclei (Hoechst 33342, blue). Scale bars in A = 10 μm, in C and D = 20 μm, in E = 40 μm. Dots in the bar graphs represent biological replicates. Data are expressed as mean ± s.d. *$P$<0.05; ** $P$<0.01, *** $P$<0.001, **** $P$<0.001; one-way ANOVA with Dunnett's multiple comparisons test.

**Data S1. (separate file)**
Differentially expressed genes in mouse sublingual glands, 7 days post-IR compared to Non-IR control.

**Data S2. (separate file)**
Differentially expressed genes in mouse sublingual glands, 14 days post-IR compared to 7 days post-IR.

**Data S3. (separate file)**
Differentially expressed genes in mouse sublingual glands, 14 days post-IR compared to Non-IR control.
